# *Pseudomonas aeruginosa* Exotoxin A is not essential for corneal disease severity or bacterial survival

**DOI:** 10.64898/2026.03.10.710780

**Authors:** James D. Begando, George R. Dubyak, Arne Rietsch, Eric Pearlman

**Affiliations:** Department of Physiology and Biophysics, University of California, Irvine, CA, USA; Department of Ophthalmology, University of California, Irvine, CA, USA; Department of Physiology & Biophysics, Case Western Reserve University, Cleveland, OH, USA; Department of Molecular Biology and Microbiology, Case Western Reserve University, Cleveland, OH, USA

## Abstract

*Pseudomonas aeruginosa* produces multiple toxins and exoenzymes that contribute to its survival and ability to cause disease. As prior studies reported an important role for type III-secretion in the severity of corneal disease in *P. aeruginosa* keratitis, we examined if there is a requirement for the type II-secreted cytotoxin Exotoxin A (ToxA) in bacterial persistence and disease severity in infected murine corneas. Using independently generated Δ*toxA* mutants and complemented strains, we found that ToxA is expressed in vivo, but ToxA deletion did not significantly affect bacterial replication, neutrophil recruitment or disease severity. These findings contrast with an earlier study identifying a critical role for ToxA.

## INTRODUCTION

Bacterial keratitis caused by *Pseudomonas aeruginosa* is a vision-threatening corneal infection characterized by rapid tissue destruction, particularly among contact lens users (1). Infection triggers epithelial injury and robust production of pro-inflammatory cytokines, driving recruitment of neutrophils and monocytes into the corneal stroma (2). Disease severity is a consequence of direct bacterial cytotoxicity and collateral host-mediated tissue damage (3). During infection, *P. aeruginosa* produces multiple virulence determinants, including proteases, phospholipases, hemolysins, type III secreted effectors (ExoU, ExoS, ExoT, ExoY), and the type II-secreted toxin Exotoxin A (ToxA) (4).

The pathogenesis of *Pseudomonas aeruginosa* keratitis is driven largely by type III secretion system (T3SS)-dependent virulence mechanisms. While we and others have reported on the important role for T3SS exoenzymes in *P. aeruginosa* keratitis (5–8), relatively little is known about the role of ToxA in corneal infection, a major virulence factor secreted through the type II secretion system (T2SS). In contrast to the T3SS, which injects effectors directly into host cells, the T2SS exports folded toxins into the extracellular environment, where they can target host tissues and contribute to disease pathogenesis. ToxA is a 66-kDa A-B toxin secreted through the T2SS and is mechanistically analogous to diphtheria toxin. The ToxA B (binding) domain attaches to the host cell surface receptor LRP1 (also known as CD91), initiating receptor-mediated endocytosis (9). Following translocation into the cytosol, the catalytic A domain ADP-ribosylates eukaryotic elongation factor-2 (eEF-2), halting protein synthesis and inducing rapid cell death (10). In this study, we sought to determine whether ToxA activity is also induced during infection and contributes to corneal disease severity.

## MATERIALS AND METHODS

### Construction of Δ*toxA* and complemented strains

An in-frame deletion of *toxA* was generated in the parent PAO1 strain by allelic exchange using the vector pEXG2 as previously described (11). Regions flanking *toxA* were amplified from genomic DNA and assembled into pEXG2, and the resulting construct was introduced into PAO1F by conjugation. Double-crossover mutants were isolated by gentamicin selection followed by sucrose counterselection, and deletion of *toxA* was confirmed by PCR.

For complementation, the *toxA* open reading frame with its native promoter was cloned into the mini-Tn7 delivery vector pUC18-miniTn7T-Gm and integrated at the chromosomal Tn7 attachment site using the Tn7 transposition system as previously described (12). The antibiotic resistance cassette was subsequently excised using FLP recombinase.

### Construction of *toxA* reporter strain

The *toxA* promoter reporter plasmid was constructed by replacing GFPmut3 in pP33-*p*_*exoS*_-GFP with a synthesized, codon-optimized superfolder GFP (GFPsf; BioBasic). The *p*_*exoS*_ promoter was subsequently replaced with the *toxA* promoter amplified from PAO1F genomic DNA. The parental pP33-*p*_*exoS*_-GFP plasmid was described previously (13).

### Growth conditions

*P. aeruginosa* strains were cultured at 37°C with shaking in dialyzed, Chelex-tryptic soy broth dialysate-free (TSB-DF) supplemented with 1% glycerol and 100 mM monosodium glutamate, adapted from previously described conditions for induction of ToxA expression and keratitis infection models (14, 15). Overnight cultures were subcultured to late-log phase (OD_600_ ≈ 0.9), washed, and resuspended in PBS prior to infection.

### Mouse corneal infection model

C57BL/6 mice were purchased from the Jackson Laboratory and bred in house. Male and female mice ages 6-12 weeks old were used for experiments. All procedures were approved by the University of California, Irvine Institutional Animal Care and Use Committee.

Overnight cultures of *P. aeruginosa* PAO1F or mutant strains were grown in TSB-DF, subcultured to late-log phase (OD_600_ ≈ 0.9), washed, and resuspended in PBS. C57BL/6 mice were anesthetized with ketamine/xylazine, the corneal epithelium was abraded with three parallel scratches using a sterile 26-gauge needle, and 2 µL of bacterial suspension (∼1 × 10^6^ CFU/eye) was applied topically as previously described (16).

### In vivo imaging and bacterial growth (CFU assay)

At 24 or 48 hours post-infection, corneas were imaged by brightfield microscopy to assess opacity or by fluorescence microscopy to detect GFP or RFP-expressing bacteria. For bacterial growth, whole eyes were homogenized in PBS using a TissueLyser II (Qiagen), serially diluted, and plated on LB agar for CFU.

### Corneal opacity and fluorescence quantification

Corneal opacity was quantified by digital image analysis using a constant circular region of interest centered on the cornea. Images were analyzed to identify opaque regions using dataset-specific pixel intensity cutoffs, glare pixels were excluded, and percent corneal opacity (opaque area) was calculated as previously described (17). GFP and RFP fluorescence were quantified from the same region of interest using integrated density measurements in ImageJ.

### Immunoblot analysis of bacterial lysates and culture supernatants

PAO1F, ΔtoxA, and complemented strains were grown overnight in TSB-DF, subcultured the following day, and grown to late-log phase (OD_600_ ≈ 0.9). Cultures were separated into cell pellet and supernatant fractions by centrifugation. Supernatants were filtered to remove bacteria, and equal culture volumes were concentrated by trichloroacetic acid (TCA) precipitation. Samples were resuspended in 1× SDS sample buffer.

Proteins were resolved on 10% SDS-PAGE gels, transferred to PVDF membranes, and probed with anti-ToxA (Sigma-Aldrich, P2318; 1:5000) and anti-RpoA (Biolegend, Cat#663104, 1:10000) in parallel. Fluorescent secondary antibodies were used for detection (anti-rabbit IRDye-800 for ToxA, anti-mouse IRDye-680 for RpoA).

### Immunoprecipitation and immunoblot analysis of corneal lysates

C57BL/6 mice were infected as described above, and corneas were harvested 24 hours post-infection. Infected corneas were pooled 4 per condition and then homogenized in denaturing lysis buffer (50 mM Tris-HCl pH 7.5, 150 mM NaCl, 1% SDS, 1 mM EDTA, protease inhibitors), heated, and clarified by centrifugation. Lysates were diluted to reduce SDS concentration and incubated overnight at 4°C with anti-ToxA antibody pre-bound to Protein G magnetic beads. Beads were washed with Tris-buffered saline followed by a high-salt wash (500 mM NaCl) to reduce nonspecific binding, and bound proteins were eluted in reducing Laemmli sample buffer.

Eluates were resolved by SDS-PAGE, transferred to PVDF membranes, and probed with anti-ToxA antibody (Sigma-Aldrich, P2318) followed by HRP-conjugated anti-rabbit secondary antibody (Cell Signaling Technology, #7074). Signal was detected by chemiluminescence.

### Flow cytometry

To recover cells, infected corneas were dissected and incubated at 37°C for 45 to 60 minutes in RPMI containing collagenase (5 mg/mL), DNase I (0.125 mg/mL), penicillin-streptomycin, non-essential amino acids, HEPES, sodium pyruvate, 10% FBS, and 10 mM CaCl_2_. Samples were gently agitated every 15 minutes to promote tissue digestion. Digestion was stopped by quenching with FBS, and cells were filtered through a 70 μm strainer, pelleted by centrifugation, and resuspended in flow cytometry buffer.

All subsequent steps were performed at 4°C. Cells were incubated with anti-CD16/32 to block Fc receptors, followed by staining with fixable viability dye eFluor780 and fluorophore-conjugated antibodies against CD45 (StarBright Violet 670), CD11b (StarBright Violet 790), Ly6G (Brilliant Violet 421), Ly6C (StarBright 575), and CCR2 (APC). Cells were fixed with paraformaldehyde, washed, and acquired on an ACEA NovoCyte flow cytometer using NovoExpress software. Leukocytes were gated as singlets and live cells, followed by CD45^+^CD11b^+^ myeloid cells. Neutrophils were identified as Ly6G^+^ and inflammatory monocytes as Ly6C^+^, CCR2^+^.

### Statistical analysis

Statistical significance was determined using one-way ANOVA followed by Dunnett’s multiple comparison test for in vivo studies. All analyses were performed using GraphPad Prism. Differences were considered significant when the P value was <0.05.

## RESULTS

### ToxA is expressed in the cornea during PAO1F infection

To assess ToxA expression in vivo, mice were infected with a *toxA* promoter reporter strain and corneas were examined at 0, 24, and 48 hours post-infection. Constitutive RFP fluorescence indicates bacterial localization within the cornea, while GFP fluorescence shows ToxA promoter activity. GFP intensity increased at 24 and 48 hours post-infection (Fig. 1A-C).

**Figure 1.**
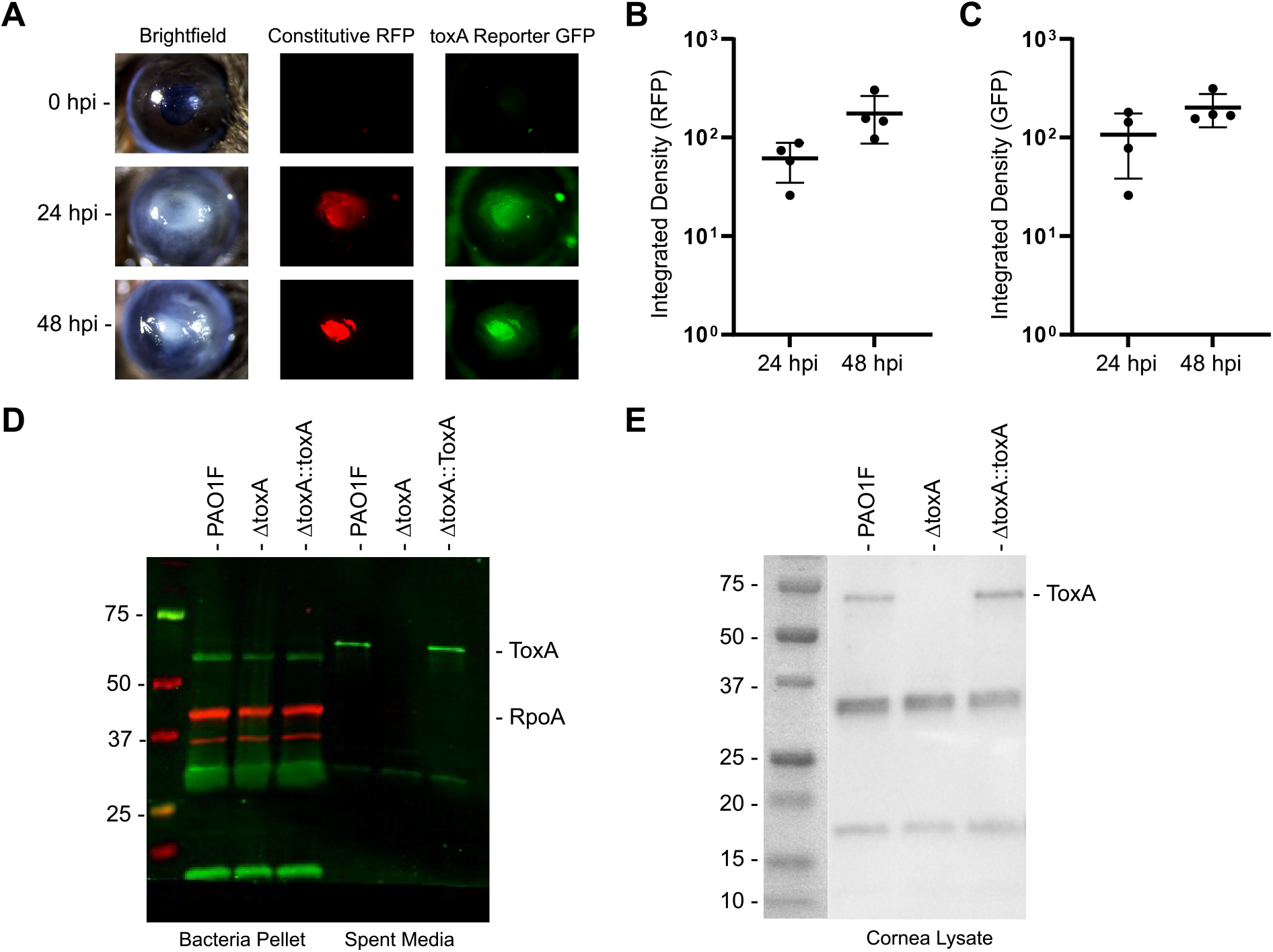
In vivo expression of ToxA during *P. aeruginosa* corneal infection. (A) Representative brightfield, constitutive RFP, and *toxA* promoter-driven GFP fluorescence images of corneas infected with a *toxA* reporter strain at 0, 24, and 48 hours post-infection. RFP indicates bacterial presence, while GFP reflects *toxA* promoter activity. (B) RFP fluorescence quantified as a measure of bacterial growth. (C) GFP fluorescence quantified as a measure of *toxA* expression. (D) Immunoblot detection of ToxA in bacterial pellet and corneal lysate samples from the indicated strains. ToxA is present in PAO1F and the complemented strain but absent in Δ*toxA*. Housekeeping RpoA protein serves as a loading control. (E) ToxA protein detected in corneal lysates by immunoblotting at the indicated time points. Each point represents a single cornea. Error bars represent mean ± SD. Statistical analysis was performed using one-way ANOVA. P>0.05 for all comparisons.

As a second approach to determine ToxA production during infection, an in-frame Δ*toxA* mutant was generated in the PAO1F background, as well as a complemented strain. Immunoblot analysis confirmed loss of ToxA protein in the Δ*toxA* mutant and restoration in the complemented strain, validating successful deletion and complementation (Fig. 1D). In infected corneas, lysates collected at 24 hours post-infection were subjected to ToxA immunoprecipitation followed by immunoblotting, which detected ToxA protein in wild-type and complemented strains but not in corneas infected with the Δ*toxA* mutant (Fig. 1E). Together, these results demonstrate that ToxA is produced in PAO1 infected corneas.

### No difference in *P. aeruginosa* growth or corneal disease severity in the absence of ToxA

We next tested whether deletion of *toxA* altered disease severity. Mice were infected with PAO1F, the Δ*toxA* mutant, or the complemented strain (Δ*toxA*::*toxA*) and evaluated 24 and 48 hours after infection. PAO1F infection produced a severe, rapidly progressive corneal infection. Brightfield imaging revealed no significant differences in mice infected with PAO1F, Δ*toxA* or Δ*toxA*::*toxA* (Fig. 2A, 2B). Similarly, there was no significant difference in GFP fluorescence (Fig. 2A, 2C). Viable bacteria, measured by CFU, were also similar at both time points in corneas infected with PAO1F, Δ*toxA*, or Δ*toxA*::*toxA* (Fig. 2D). Together, these data indicate that loss of *toxA* does not regulate bacterial growth or corneal disease severity in this model.

**Figure 2.**
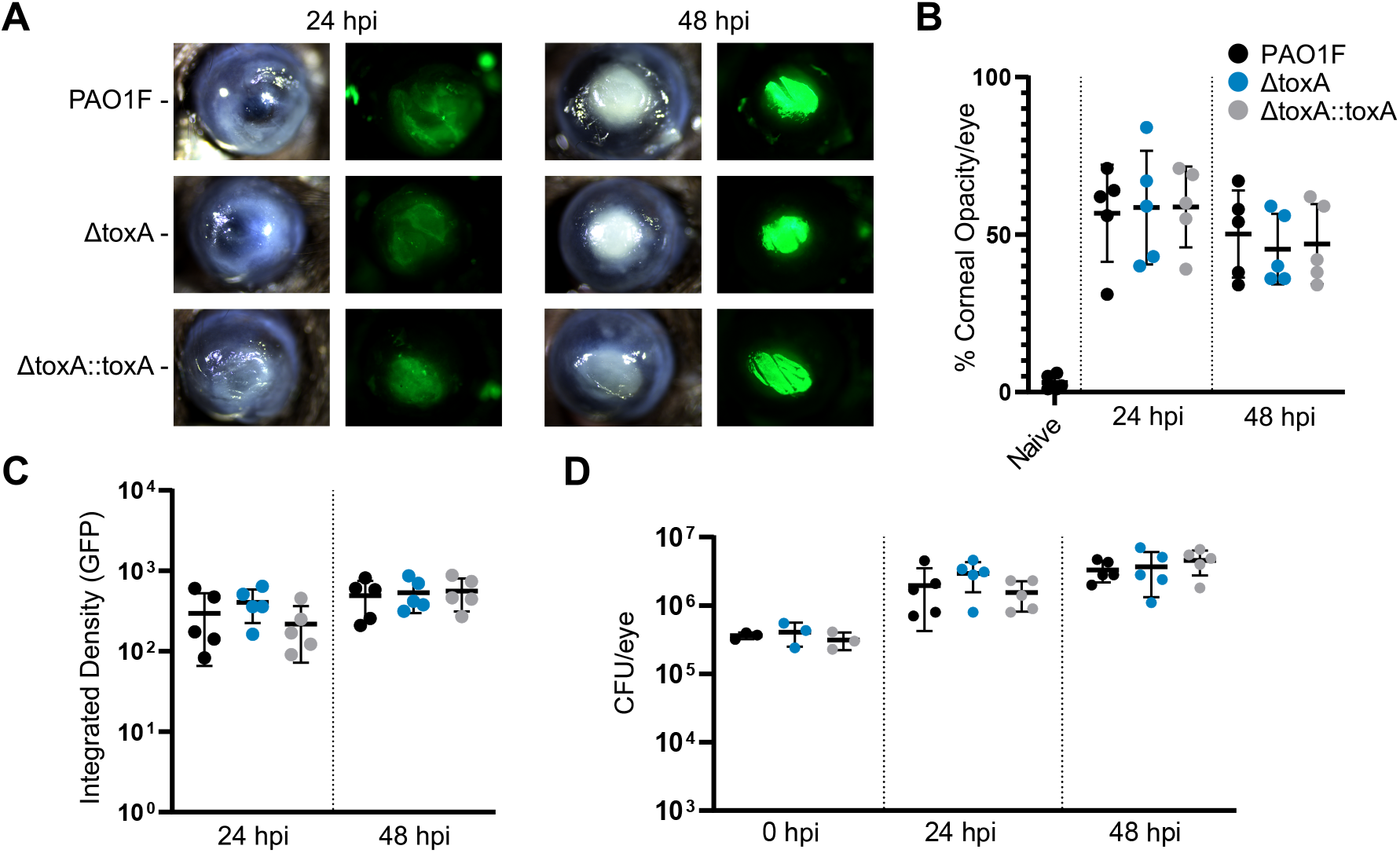
Corneal infection with PAO1F ToxA mutants. **(A)** Representative brightfield and constitutive GFP fluorescence images of infected corneas at 24 and 48 hours post-infection. **(B)** Corneal opacity quantified and expressed as a percentage. **(C)** GFP fluorescence quantified as a measure of bacterial growth. **(D)** Viable bacterial growth measured by colony-forming unit (CFU) from homogenized corneas. Each point represents a single cornea. Error bars represent mean ± SD. Statistical analysis was performed using one-way ANOVA. P>0.05 for all comparisons.

### Host inflammatory cell recruitment is not altered by ToxA deletion

To assess leukocyte infiltration, we performed flow cytometry at 24 hours post-infection and found that approximately 90% of infiltrating CD45^+^CD11b^+^ cells were neutrophils and ∼10% were monocytes (Fig. 3A), consistent with our findings in infected patients and prior work using this keratitis model (16, 18). Total neutrophil and monocyte numbers did not differ significantly between groups (Fig. 3B, 3C). After 48 hours, ∼90% of monocytes expressed CCR2, consistent with an inflammatory monocyte phenotype (16), and the proportion of CCR2^+^ monocytes was not significantly different between PAO1, Δ*toxA*, and the complemented strain (Fig. 3D). Overall, inflammatory cell recruitment was not affected in the absence of ToxA.

**Figure 3.**
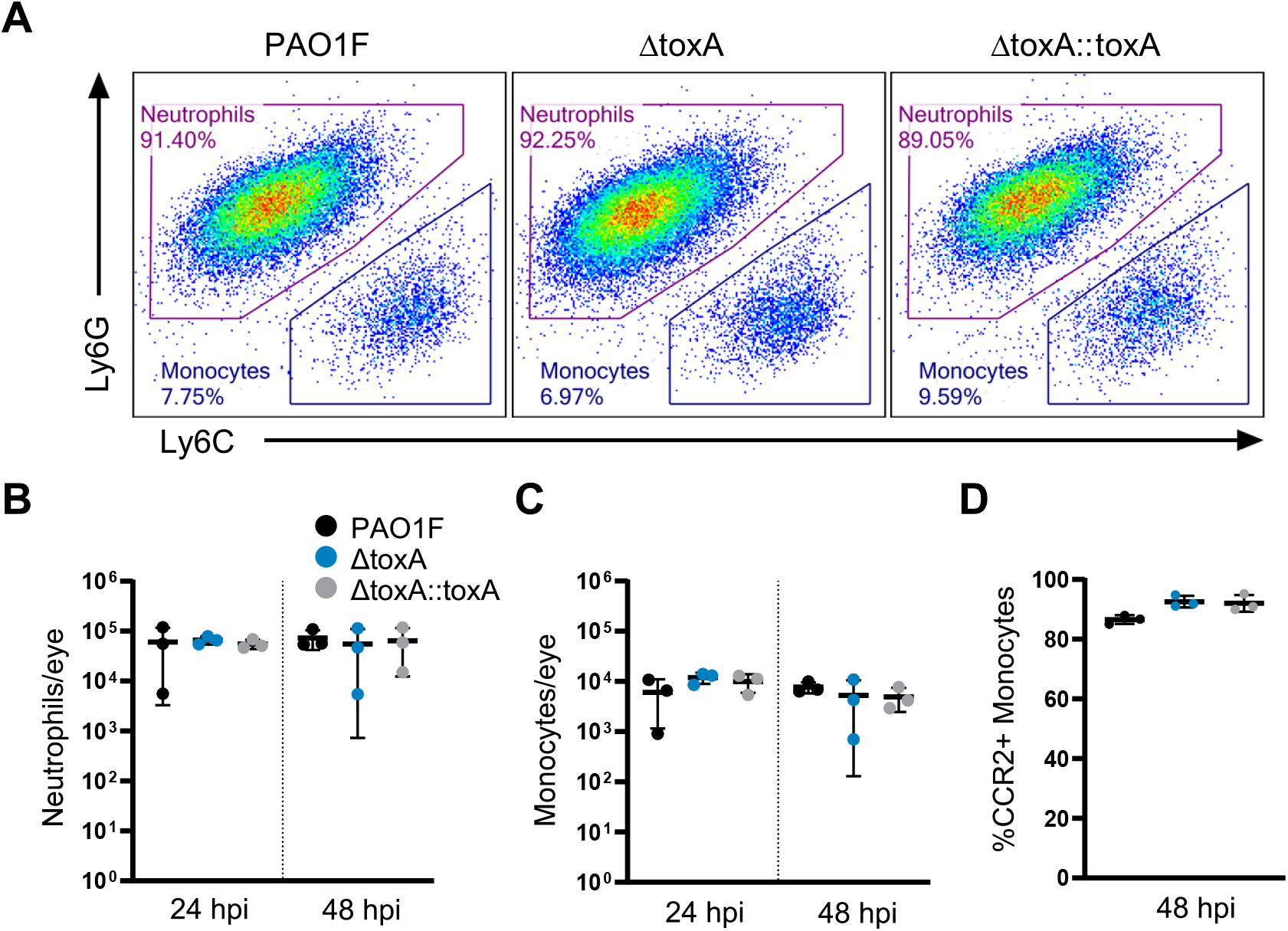
Neutrophil and monocyte recruitment to infected corneas. **(A)** Representative flow cytometry plots from collagenase-digested corneas at 24 hours post-infection (hpi) with PAO1F, Δ*toxA*, or complemented strains of *P. aeruginosa*. Cells were stained for Ly6G and Ly6C to identify neutrophils (Ly6G^+^) and monocytes (Ly6C^+^). Percentages represent frequencies within the CD45^+^/CD11b^+^ population. **(B)** Total number of neutrophils per cornea at 24 and 48 hpi. **(C)** Total number of monocytes per cornea at 24 and 48 hpi. **(D)** Percentage of monocytes expressing CCR2 at 48 hpi. Each point represents a single cornea. Error bars represent mean ± SD. Statistical analysis was performed using one-way ANOVA. P>0.05 for all comparisons.

### ToxA is not required for corneal disease in strains with reduced or no T3SS production

To determine whether type III secretion system (T3SS) activity masked a contribution of ToxA, we repeated these experiments in strains with reduced or absent T3SS function. Because the PAO1F background exhibits high T3SS activity that can dominate corneal pathology, we asked whether lowering or eliminating T3SS would reveal a ToxA-dependent phenotype. The T3SS-low strain used here is a PAO1 background that exhibits reduced and delayed type III secretion gene expression, producing weaker and slower T3SS-dependent phenotypes in vitro (19). The T3SS-low strain showed reduced disease at 24 hours post-infection but recovered by 48 hours post-infection. However, within this background, deletion of *toxA* did not alter corneal opacity, GFP signal, or bacterial growth (Fig. 4A-D). Similarly, the Δ*pscD* strain showed that absence of Type III secretion reduced infection overall, yet *toxA* deletion again had no effect on pathology or CFU (Fig. 4E–H). Across both backgrounds, the absence of ToxA had no discernable effect on corneal opacity or bacterial survival.

**Figure 4.**
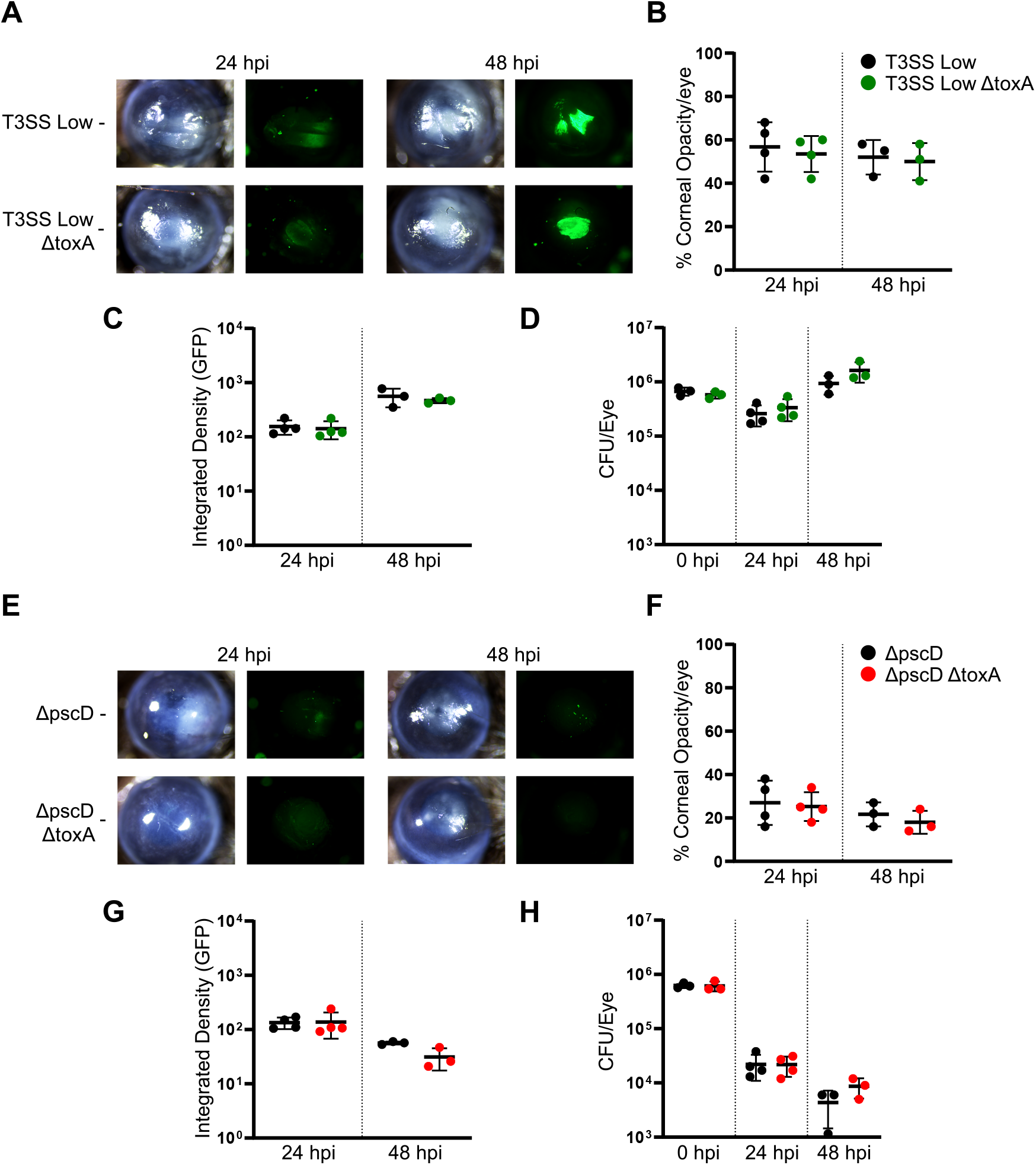
The effect of ToxA in T3SS low expressing and a T3SS-deficient (ΔpscD) strain. **(A–D)** Mice were infected with T3SS Low PAO1 or T3SS Low Δ*toxA* strains and analyzed at 24 and 48 hours post-infection (hpi). **(A)** Representative brightfield and constitutive GFP fluorescence images of infected corneas. **(B)** Corneal opacity quantified and expressed as a percentage. **(C)** GFP fluorescence quantified as a measure of bacterial growth. **(D)** Viable bacterial growth measured by colony-forming units (CFU) from homogenized corneas. **(E–H)** Mice were infected with Δ*pscD* or Δ*pscD* Δ*toxA* strains and analyzed at 24 and 48 hours post-infection (hpi). **(E)** Representative brightfield and constitutive GFP fluorescence images of infected corneas. **(F)** Corneal opacity quantified and expressed as a percentage. **(G)** GFP fluorescence quantified as a measure of bacterial growth. **(H)** Viable bacterial growth measured by colony-forming units (CFU) from homogenized corneas. Each point represents a single cornea. Error bars represent mean ± SD. Statistical analysis was performed using one-way ANOVA. P>0.05 for all comparisons.

## DISCUSSION

We and others reported a role for T3SS in the pathogenesis of *P. aeruginosa* keratitis (5– 8). We focused primarily on the ExoS-expressing strain PAO1 rather than ExoU-producing strains, as most clinical isolates express either ExoS or the ExoU phospholipase, but rarely both (20). Our earlier findings identified a role for ExoS and ExoT ADP ribosyl transferase (ADPRT) in inhibiting NADPH oxidase assembly and ROS production. We also demonstrated that ExoS ADPRT regulates inflammasome usage in neutrophils (6).

Although these studies highlight the importance of T3SS-dependent virulence mechanisms in *P. aeruginosa* keratitis, the contribution of other secreted toxins to disease progression remains less clear. To address this question, we investigated the contribution of the ADP-ribosylating cytotoxin Exotoxin A (ToxA), which is secreted through the type II secretion system (T2SS). The role of ToxA in *P. aeruginosa* keratitis has largely been defined by a single foundational report by Pillar and Hobden, who showed that Δ*toxA* mutants were rapidly cleared from the cornea and resulted in reduced disease severity compared with wild-type strains (14). Using isogenic mutants generated in both PAO1 and 19660 (ExoU expressing) backgrounds, they demonstrated that ToxA was not required for bacterial adherence to wounded corneal epithelium or for initiation of infection, but was critical for bacterial persistence, sustained inflammation, and progression to severe disease. Over the past two decades, this study has been cited extensively, and the prevailing view is that ToxA plays a major role in *P. aeruginosa* keratitis.

Given these prior findings, we generated toxA deletion and complemented strains of PAO1 in the presence or absence of T3SS activity and examined their virulence in murine corneas. Although we confirmed that *toxA* is expressed in infected corneas during infection, we observed no differences in bacterial growth, disease severity, or inflammatory cell recruitment between PAO1 and Δ*toxA* strains. Despite matching the inoculum, scarification method, mouse strain and age, and similar bacterial growth conditions reported by Pillar and Hobden (14), Δ*toxA* mutants did not show reduced virulence, yielding results that differ from this earlier report. Our findings indicate that ToxA is not required for disease progression under the conditions tested in this model.

A limitation of the present study is that we evaluated ToxA function only in an ExoS-producing PAO1 background, whereas Pillar and Hobden examined both PAO1 and strain 19660, an ExoU-producing lineage (14). ExoU strains that produce a highly cytotoxic phospholipase typically cause more rapid and severe corneal disease than ExoS strains (18). However, ExoS remains a major determinant of keratitis pathogenesis, promoting neutrophil apoptosis and enhancing bacterial survival in the cornea (21, 22). The absence of a detectable role for ToxA, even in a T3SS low strain, suggests that ToxA is not required for disease when type III secretion-dependent virulence mechanisms are active. Because ExoU strains are even more virulent, their presence would be expected to further obscure any contribution from secondary toxins such as ToxA.

A potential explanation for the absence of a detectable ToxA phenotype in this model may lie in species-specific differences in inflammasome activation pathways. A recent study showed that eEF-2-targeting toxins, including ToxA, activate the NLRP1 inflammasome and disrupt epithelial barrier integrity in human cells (23), supporting the possibility that ToxA may engage NLRP1-dependent inflammatory pathways. Mechanistically, eEF-2 inhibition induces translational arrest and ribosome stalling, activating the ZAKα-dependent ribotoxic stress response and downstream p38 signaling, which in turn promotes NLRP1 activation through phosphorylation of a primate-specific N-terminal regulatory region (24–27). Through this mechanism, ZAKα-activating ribotoxins such as anisomycin can activate human NLRP1, but not murine NLRP1B as anisomycin fails to induce NLRP1-dependent IL-1β release in murine keratinocytes and macrophages (27). In contrast, murine NLRP1B is activated primarily by direct proteolytic cleavage of the receptor, as demonstrated for anthrax lethal factor (28). Similarly, viral proteases also activate NLRP1 through cleavage, but the sites and requirements differ between species, indicating that human and murine NLRP1 respond to distinct upstream signals (25).

In the cornea, disease severity caused by *P. aeruginosa* is driven largely by recruited neutrophils in infected patients (3, 16, 18). Inflammasome activation by ribotoxic stress in human neutrophils may represent an important mechanism by which ToxA contributes to disease. Determining whether this pathway contributes to corneal pathology will require models that more accurately reflect human inflammasome signaling.

Taken together, these findings indicate that ToxA is not a required determinant of corneal pathology in mice and highlight the importance of evaluating ToxA function in human infection models.

## Author contributions

JB and AR performed experiments and generated graphs and images. AR generated all the mutants and complemented strains.

JB, GD, AR, and EP designed the experiments and wrote the manuscript.

